# High-resolution cryo-electron microscopy structure of the *Escherichia coli* 50S subunit and validation of nucleotide modifications

**DOI:** 10.1101/695429

**Authors:** Vanja Stojković, Alexander G. Myasnikov, Iris D. Young, Adam Frost, James S. Fraser, Danica Galonić Fujimori

## Abstract

Post-transcriptional ribosomal RNA (rRNA) modifications are present in all organisms, but their exact functional roles and positions are yet to be fully characterized. Modified nucleotides have been implicated in the stabilization of RNA structure and regulation of ribosome biogenesis and protein synthesis. In some instances, rRNA modifications can confer antibiotic resistance. High-resolution ribosome structures are thus necessary for precise determination of modified nucleotides’ positions, a task that has previously been accomplished by X-ray crystallography. Here we present a cryo-electron microscopy (cryo-EM) structure of *Escherichia coli* (*E. coli*) 50S subunit at an average resolution of 2.2Å as an additional approach for mapping modification sites. Our structure confirms known modifications present in 23S rRNA and additionally allows for localization of Mg^2+^ ions and their coordinated water molecules. Using our cryo-EM structure as a testbed, we developed a program for identification of post-transcriptional rRNA modifications using a cryo-EM map. This program can be easily used on any RNA-containing cryo-EM structure, and an associated Coot plugin allows for visualization of validated modifications, making it highly accessible.

## INTRODUCTION

Ribosomes are complex cellular machines responsible for protein synthesis. Accurate translation by the ribosome requires specific post-transcriptional and post-translational modifications of ribosomal RNA (rRNA) and ribosomal proteins, respectively. The roles of individual rRNA modifications are still poorly understood, and while some modulate the function of the ribosome, others are important in RNA folding and stability (1–4). The majority of rRNA modifications are located in functional regions of the ribosome: the decoding center, the peptidyl transferase center (PTC), the exit tunnel, or the interface between ribosomal subunits. In *E. coli*, not a single rRNA modification is essential for the survival of the cell. However, the presence of individual modifications confers advantages under certain growth or stress conditions (3). For instance, modifications of rRNA have emerged as one of the most clinically relevant mechanisms of resistance to ribosome-targeting antibiotics (5–8). Differential expression and post-translational modifications of ribosomal proteins, as well as differences in rRNA modifications, can lead to heterogeneity in ribosome composition which can result in ‘specialized ribosomes’. These ribosomes can have a substantial impact on the relative abundances of proteins being produced and consequently influence an organism’s adaptability to a variety of environmental factors (9–11). For example, in bacteria, ribosome heterogeneity has been identified as a mechanism for stress adaptation (12).

Visualizing rRNA modifications is important for understanding their impact on the translating ribosome, and can aid in the development of novel antibiotics. Recently, rRNA and ribosomal protein modifications have been visualized in several high-resolution X-ray and cryo-electron microscopy (cryo-EM) structures of bacterial ribosome (2.3Å resolution) (13), including *E. coli* ribosome (2.4Å) (14), parasitic (2.8Å - cryo-EM) (15) and human ribosomes (2.8Å - cryo-EM) (16, 17). In certain instances, assignments of nucleotide modifications have been subsequently revised in the light of improved data (18), highlighting the challenges in assigning nucleotide modifications in near-atomic resolution structures.

Here, we report high-resolution cryo-EM structure of the *E. coli* 50S subunit and validate rRNA modifications using a program we developed for quantifying post-transcriptional modifications, which we named qPTxM. This program probes the map density at each nucleotide to identify all possible modifications. The modifications found to be well-supported by the map are written to a copy of the model and can be easily examined in Coot using a custom plugin (19). Additionally, we confirmed the assigned nucleotide conformations using the X3DNA software system (20).

## MATERIAL AND METHODS

### *E. coli* 50S ribosomal subunit purification

50S ribosomal subunit was purified from *E. coli* MRE600 strain using modified version of previously published protocol (21). In short, cells were grown to an OD_600_ of 0.5 in LB media at 37°C and 220 rpm. Cells were pelleted by centrifugation, washed with wash buffer (20 mM Tris pH 7.5, 100 mM NaCl, 10 mM MgCl_2_) and stored at −80°C. The pellet corresponding to 750 mL of growth was resuspended in Buffer A (20 mM Tris pH 7.5, 300 mM NH_4_Cl, 10 mM MgCl_2_, 0.5 mM EDTA, 6 mM β-mercaptoethanol and 10 U/mL of SuperASE-In (Ambion)). PMSF was added to a final concentration of 0.1 mM. The cell resuspension was lysed using microfluidizer at 10,000 psi with three passages. The lysate was centrifuged at 16,100 rpm, (Beckman Ti45 rotor) for 30 min, at 4°C, to remove cell debris. Without disturbing the pellet, supernatant was removed and centrifuged for additional 30 min. The resulting supernatant was loaded onto a 32% sucrose cushion prepared in Buffer A and centrifuged at 28,000 rpm (Beckman SW41Ti rotor) for 16 h, at 4°C, to obtain the crude 50S pellet. This pellet was slowly resuspended at 4°C in Buffer A to homogeneity. Any presence of non-resuspended particles was removed by a short centrifugation at 10,000 rpm, for 10 min at 4°C.

Crude 50S ribosomal subunits were further purified on 10-40% sucrose gradients. Sucrose gradients were prepared in Buffer A using BioComp Gradient Master. Approximately 650 pmol of crude 50S subunits were loaded on each gradient and centrifuged at 28,000 rpm (Beckman SW41Ti rotor) for 16 h, at 4°C. Fractions were collected from top to bottom using BioComp Piston Gradient Fractionator. The sample absorbance was recorded using UV reader and the peak corresponding to 50S was pooled for precipitation with PEG 20,000. A final concentration of 10.5% PEG 20,000 was slowly added to the pooled fractions, incubated on ice for 10 min and centrifuged at 10,000 rpm, for 10 min at 4 °C. The pure 50S pellet was dissolved in Buffer A and filtered using 0.22 µm low-binding Durapore PVDF filters (Millipore).

### Cryo-EM analysis

Purified 50S ribosomal subunits were diluted from 2 mg/mL to 0.5 mg/mL in Buffer A, applied to 300-mesh carbon coated (2nm thickness) holey carbon Quantifoil 1.2/1.3 grids (Quantifoil Micro Tools) and flash-frozen as described in ref (22). Data were collected on the in-house Titan Krios X-FEG instrument (Thermo Fisher Scientific) operating at an acceleration voltage of 300 kV and a nominal underfocus of Δz = −0.2 to −1.5 μm at a magnification of 29.000 (nominal pixel size of 0.822Å). We recorded 1889 movies using a K2 direct electron detector camera in super-resolution mode with the total dose of 80e fractionated over 80 frames (80 individual frames were collected, starting from the 1^st^ one). Total exposure time was 8 s. Images in the stack were aligned using MotionCor2 (23). Before image processing all micrographs were checked for the quality and 1610 best based on the power spectrum quality and particle distribution, were selected for the next step of image processing. The contrast transfer function of each image was determined using GCTF (24) as a standalone program. For particle selection we have used Relion3 autopicking procedure which picked 193,000 particles (25). For the first steps of image processing we used coarsened data binned by a factor of 8 (C8 images) and 60 classes. First, we applied 2D classification to remove only images with ice or other contaminants (162,746 particles left), followed by 3D classification with bin by 4 data (C4) to remove bad particles (141,549 good particles left). After one round of 3D refinement with C2 data we performed additional round of 3D classification with high T (from 10 to 20) in order to select the best particles. This was followed by 3D refinement with C1 data. For the initial refinement we used spherical mask, which was followed by further refinement using mask around stable part of 50S (excluding L1 stalk, L7/L12 region). A further improved cryo-EM map was obtained by using CTF-refinement procedure from Relion3. The Post-processing procedure implemented in Relion3 (25) was applied to the final maps with appropriate masking, B-factor sharpening (automatic B-factor estimation was −55.85) and resolution estimation to avoid over-fitting (Final Resolution after Post-processing with 50S mask applied was 2.68Å). Subsequently the stack of CTF-refined particles was processed in a new version of CryoSPARC v2.0 (26). After homogeneous refinement resolution was improved to 2.5Å. The same stack of particles was additionally refined in cisTEM (27). After Auto-Refine we attained 2.4Å resolution and with further per particle CTF refinement the resolution was improved to 2.2Å (Figure 1 and Supplementary Figure S1). This map after Sharpen3D (27) was used for model building and map interpretation.

**Figure 1.**
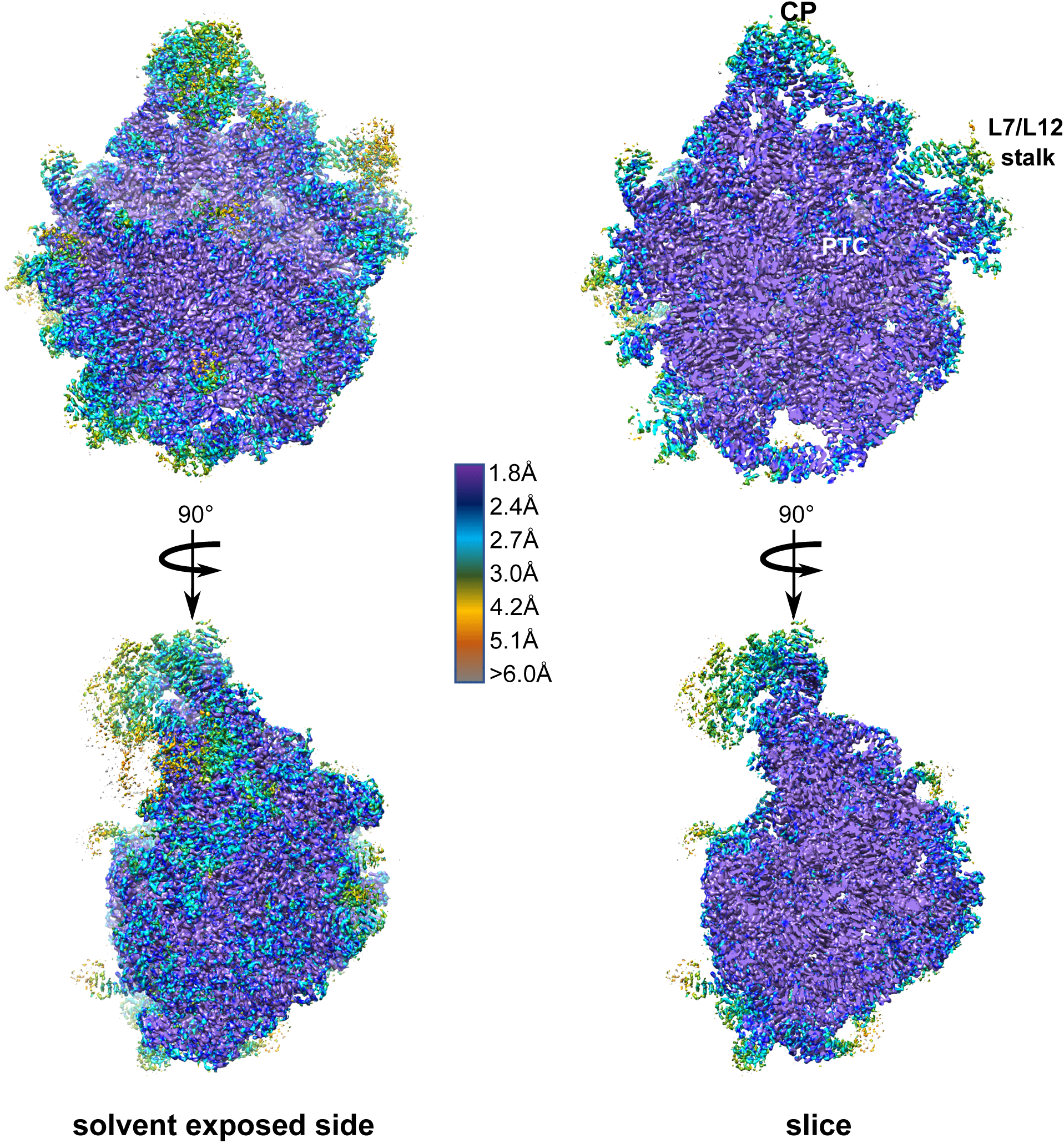
3D cryo-EM map of the 50S *E. coli* subunit colored according to local resolution. Left, solvent exposed side view; right, cut-through view. Features in the 50S subunit include the central protuberance (CP), peptidyl-transferase center (PTC), L7-L12 region (L7/12 stalk).

### Atomic model building and refinement

The final model of the 50S subunit was generated by multiple rounds of model building in Coot (19) and subsequent refinement in PHENIX (28). Modified nucleotides were drawn and the restraints for the atomic model fitting and refinements were generated using JLigand (29). The atomic model of the 50S subunit from the *E. coli* ribosome structure (PDB 4YBB) (14) was used as a starting point and refined against the experimental cryo-EM map by iterative manual model building and restrained parameter-refinement protocol (real-space refinement, positional refinement, and simulated annealing). Final atomic model comprised ∼154836 atoms (excluding hydrogens) across the 3016 nucleotides and 3356 amino acids of 29 ribosomal proteins. Proteins L3, L10 and L31 were not modelled in. In addition, 185 Mg^2+^, 3897 water molecules, one Zn^2+^, and one Na^+^ were included in the final model. Prior to running MolProbity (30) analysis, nucleotides 878-898, 1053-1107, 2099-2188 of 23S rRNA, and ribosomal proteins L9 and L11 were removed, due to their high degree of disorder. Protein residues show well-refined geometrical parameters (allowed regions 4.68%, favored regions 95.26%, and 0.07% outliers in Ramachandran plots; Supplementary Table S1). Figures were prepared using UCSF Chimera 1.13 (31).

### qPTxM program for quantitative identification of post-transcriptional modifications: development and analysis

qPTxM was developed for detection of post-transcriptional modifications based on evidence from an X-ray or EM map. It requires a map, an atomic model refined into it, and the estimated average resolution of the map. First, qPTxM reads the model and constructs a copy containing only the major conformer of each residue or nucleotide, removes all hydrogens, and removes any post-transcriptional modifications already modeled. The resulting model is used to generate a calculated map. Using the calculated and experimental maps, a difference map is generated, where positive difference density features will indicate sites of possible unmodeled atoms. qPTxM then examines each nucleotide and corresponding density in the experimental and difference maps for possible modifications (Figure 2). First, the correlation coefficient between the calculated and experimental maps is calculated using 8-point interpolation at the position of each atom in the unmodified nucleotide. If the correlation coefficient is worse than 0.6, the quality of the density is considered insufficient to support the presence of modifications, and no modifications will be suggested at this nucleotide. The default threshold of 0.6 is a lower bound that may be adjusted upward for higher confidence and lower sensitivity where high resolution data permit more stringent criteria.

**Figure 2.**
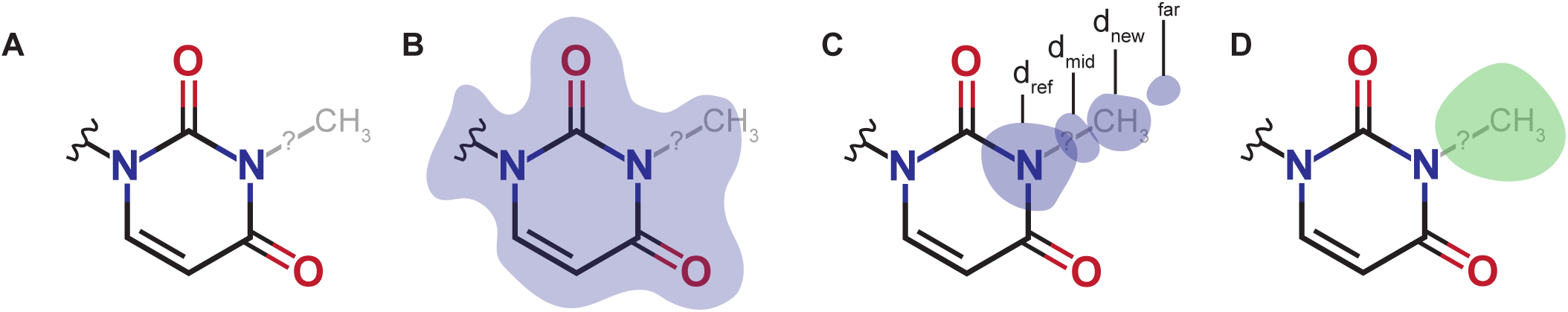
Methodology for assignment of post-transcriptional modifications (PTxMs). (**A**) For each possible modification, the new atoms are modeled with ideal geometry on a copy of the model. (**B**) Correlation of the model with the map is used to remove modifications for which the model is already a poor fit in the density. (**C**) Ratios of densities at several locations on the modified residue are used to assess whether the density profile along the vector of each new covalent bond matches expectations. By default, d_far_ may be no more than twice d_new_, d_far_ may not be more than d_mid_, and d_ref_ may be no more than three times d_new_. (**D**) A difference map is calculated between the experimental map and a map calculated from the (unmodified) model, and only the modifications with the strongest experimental and difference map densities at the new atom positions (by default, the 50% strongest by each measure) are considered. An illustration of these parameters is shown for N3-methyluridine.

Second, densities at four groups of positions in the experimental map are calculated and compared. For each modification we define one or more reference atoms (ref), one or more new atoms (new), positions midway between the reference and the new atoms (mid), and positions half a covalent bond beyond the new atoms along the direction of the bond (far). We obtain the experimental map densities at each position by 8-point interpolation and calculate the average density within each of these categories. Ratios of these densities d_ref_, d_mid_, d_new_ and d_far_ are used as indications of how closely the shape of the density around the modification matches our expectation, and we specifically select three comparisons that help eliminate a large number of cases where features in the map at the proposed modification position can be explained by water molecules, ions, and protrusions from other parts of the protein or nucleic acid model. We have found that requiring d_far_ ≤ 1.5•d_new_, d_far_ ≤ d_mid_ and d_ref_ ≤ 3•d_new_ filters out most of these cases. These multipliers may also be adjusted with optional arguments to qPTxM.

Third, the collection of possible modifications is sorted by d_ref_ and d_new_diff_, where the latter quantity is the density measured by 8-point interpolation at the new atom positions in the difference map, and the modifications in the lower half of d_ref_ and the lower 90% of d_new_diff_ densities are excluded from consideration. This step improves selectivity for nucleotides that have strong densities and strong evidence for additional unmodeled atoms. At all stages, proposed modifications are marked as rejected but are not removed from memory, so that all filters operate on the full set of proposed modifications. The full set is written to all_tested_ptms.out to facilitate rapid filter-only runs of qPTxM, while ptms.out contains all accepted modifications.

Finally, modifications are scored by taking the ratio d_new_/d_ref_ and inflating it by a factor between 1.2 for atoms close to the backbone (in modifications such as 7-methylguanosine (m^7^G) and 3 for atoms distant from the backbone (in modifications such as 2-methylguanosine (m^2^G)). Thus, the score is normalized depending on how far the modified position is from the backbone and whether the reference atoms are part of the aromatic ring or protrude from it. This scaling factor helps account for the steeper dropoff in density at positions further from other heavy atoms. Scores are compared against a default threshold of 0.5, which can again be adjusted with an optional argument.

Modifications passing all tests are built into a copy of the model for visualization, with the possibility of multiple modified residues at a single position, which is written to a file ptms.pdb. qPTxM also generates a plain text file ptms.out, a script goto_ptms.py, and pdfs of histograms of the correlation coefficients, scores and densities. It is possible to rerun just the filtering steps of qPTxM to regenerate plots and output (.out) files for rapid fine-tuning of the optional parameters using the argument adjust_filters_only=True, since these analyses can be run on the ptms.out file alone without repeating each of the calculations. This can be useful for adjusting sensitivity until a reasonable number of modifications pass all tests. To generate the custom script and the copy of the model with modifications in place, qPTxM should be run again without this argument once the user has settled on final parameters. The command “coot goto_ptms.py pruned.pdb ptms.pdb” will load two copies of the model in the Coot model viewing program, where pruned.pdb contains no modifications and ptms.pdb contains those passing all tests (Supplementary Figure S2). The script goto_ptms.py adds functionality to jump directly to each modification for visual inspection.

We used synthetic data to tune the behavior of qPTxM under ideal (noise-free) conditions with perfect confidence in which modifications should be discovered. We are including the tools we used to generate the synthetic data in qPTxM for the benefit of other users and developers, accessed by supplying the command line argument synthetic_data=True. Beginning with a refined map and model, we strip off any modifications already present, any alternate conformers, and all hydrogens, just as for a model to be tested. This model is then modified at random at 10% of all recognized nucleotides (*i*.*e*. all standard, unmodified nucleotides as well as any modified nucleotides the program recognizes how to strip modifications off of). Then, instead of using the experimental map supplied, the modified model is used to generate calculated structure factors (substituting for a noise-free experimental map) and the experimental map supplied is used only to inform the dimensions of the calculated map. Subsequently, qPTxM proceeds through all other steps as described above, and finishes by reporting on its accuracy. The list of (true) modified sites in the synthetic data is also written to a file synthetic_ptms.out and can be examined independently.

## RESULTS

Using a Titan Krios X-FEG instrument equipped with a K2 direct electron detector camera operated in super-resolution mode and performing reconstruction refinement in Relion3, CryoSPARC and cisTEM software, we obtained a structure of the *E. coli* 50S subunit at average resolution of 2.2Å (Supplementary Table S1). In order to avoid any model bias, the initial *ab initio* structure was generated in CryoSPARC and then used as a reference in Relion3. To achieve higher resolution, later steps of 3D refinement included only the well-ordered regions of the 50S subunit, and excluded the L1 stalk and L7/L12 region. An improved cryo-EM map was subsequently obtained using the CTF-refinement function of Relion3 and further improved using per-particle CTF refinement in cisTEM (see Methods). The resulting local resolution map is uniformly sampled, with a large portion of the map showing resolution of ∼2Å (Figure 1). Highly dynamic peripheral regions of the 50S subunit, encompassing the L1 arm, the stalk proteins L10, L11 and L7/L12 region, ribosomal proteins L9 and L31, and sections of the GTPase center, however, are disordered. While proteins L1, L7, L10 and L31 are not built in our model, the amino-terminal domains of the proteins L9 and L11 are structurally well-defined and thus these two proteins are built into the final model. Conformations and complete models of proteins L9 and L11 were modeled based on the high-resolution X-ray crystal structure (14). Overall, our structure is the highest resolution cryo-EM ribosome structure to date, allowing us to unambiguously visualize many rRNA modifications, Mg^2+^ ions and coordinated waters.

### Ribosomal modifications

Most ribosomal modifications in the 50S subunit are clustered in the PTC region and the nascent peptide exit tunnel (NPET). Excluding pseudouridines (Ψ), we modeled 23 out of 25 known post-transcriptional modifications in the 23S rRNA, including the non-planar base of dihydrouridine (D) at position 2449 (Figure 3, Supplementary Figure S3 and Supplementary Table S2). The only modifications not observed are substoichiometric thionation of cytidine 2501 (s^2^C) and disordered methylated pseudouridine 1915 (m^3^Ψ1915). Nucleotide 1915 is located at the interface between the large and the small ribosomal subunits and it has been proposed to strengthen the intersubunit contact (13). Seven out of ten known pseudouridines were well-ordered and thus are modelled in our structure (Supplementary Figure S3). Unlike uridines, pseudouridines contain an extra hydrogen-bond donor in the form of an N1 imino group, and water-mediated contact between the pseudouridine N1 imino group and the rRNA phosphate backbone is taken as an indication of pseudouridine presence. This water-mediated contact was observed for several pseudouridines (Supplementary Figure S3 and Supplementary Table S3).

**Figure 3.**
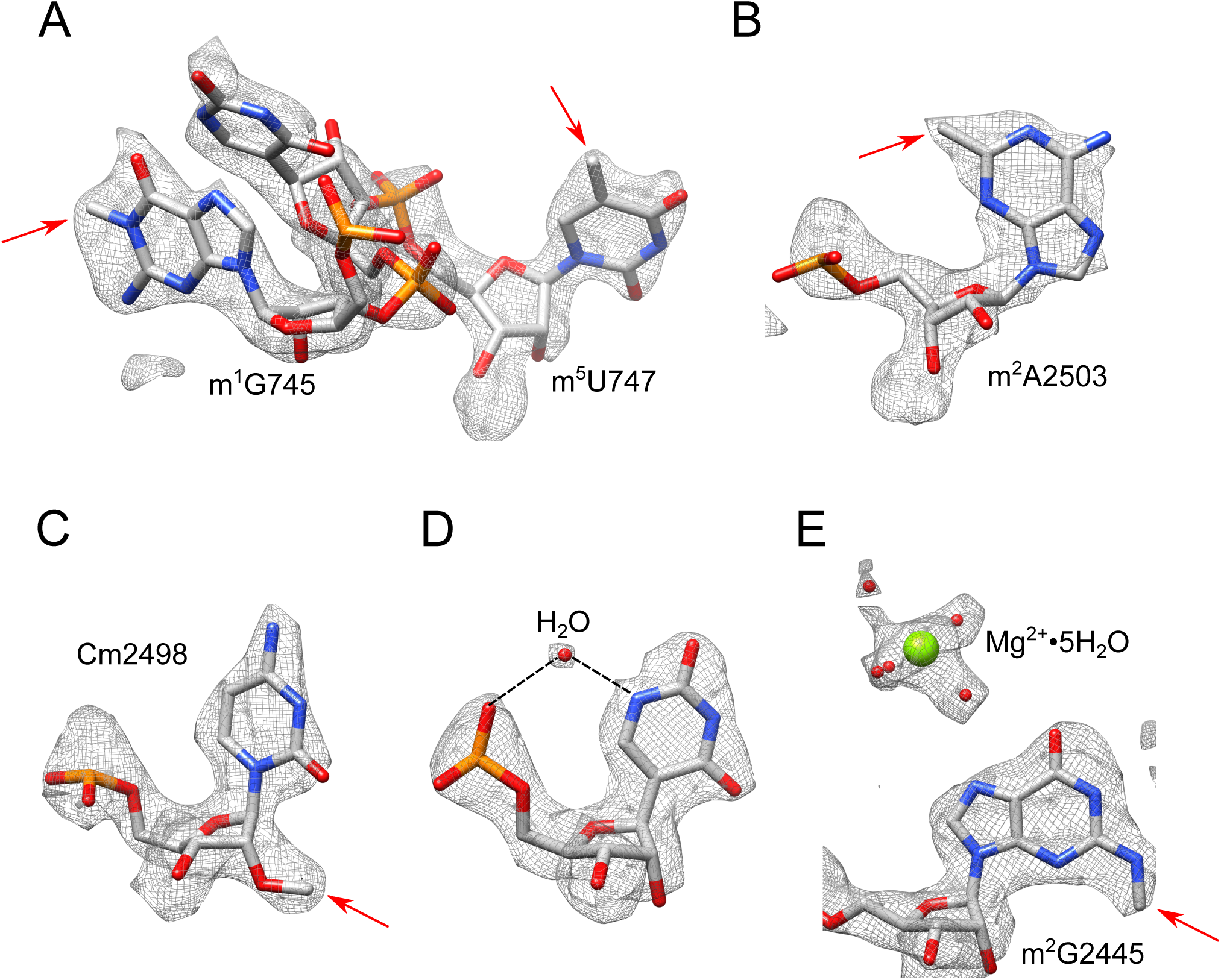
CryoEM density map allows modeling of rRNA modifications of the *E. coli* 50S ribosomal subunit. (**A-D**) CryoEM density map of selected modified nucleotides present in 23S rRNA. The modifications shown are: (**A**) 1-methylguanosine (m^1^G) 745, pseudouridine (Ψ) 746, 5-methyluridine (m^5^U) 747, (**B**) 2-methyladenosine (m^2^A) 2503, (**C**) 2’-O-methylcytidine (Cm) 2498, (**D**) pseudouridine (Ψ) 955, and (**E**) 2-methylguanosine (m^2^G). (**D, E**) Coordinated waters and Mg^2+^ are also shown.

### Validating the ribosomal modifications using density sampling: qPTxM

While there is precedent for modeling nucleotide modifications in high-resolution structures (<2.5 Å), it is important to determine the reliability of these assignments. To independently examine assigned modifications in the cryo-EM map, we developed a new program, qPTxM (for <u>q</u>uantifying <u>p</u>ost-<u>t</u>ranscriptional <u>m</u>odifications). Development of this tool was motivated by the idea that modifications can be identified directly from high-resolution maps (e.g. (17)), including the one produced here (Figure 3). The premise of qPTxM is similar to EMRinger (32), which quantifies agreement between the protein backbone and EM density maps. In both programs, the part of the structure that can be most confidently built (the nucleobase or the protein backbone) is used to assess the strength of the EM density support at positions that are predicted, with ideal geometry, to be occupied by features that require high resolution data to model accurately (the nucleotide modification or the gamma atom of the protein side chain). These approaches leverage prior knowledge of ideal geometry to objectively quantify the strength of density at positions where a model might be extended, in this case identifying possible sites of post-transcriptional modification. qPTxM produces a ranked list of candidate sites with support for interactive viewing in Coot (19) (Supplementary Figure S2) (see Methods).

To test evidence for modifications on an EM model and map, the model is first stripped of all known modifications. Since the model is invariant to the map in EM, this serves to blind the program to any modifications that may have been previously identified and to produce a *calculated map* from this model. We then compare the *calculated map* against the *experimental map*, to produce a *difference map*. Naively, we expect the *difference map* to contain difference density at the sites of unmodeled features. qPTxM examines each nucleotide and constructs a copy containing each possible modification, placing new atoms with ideal geometry. We examine the evidence for each of these modified nucleotides with a series of tests. Each of the filters described below is tunable, allowing the user to match the parameters to the quality of the map (Supplementary Table S4). We calibrated the default parameters based on this ribosome map and synthetic data. Our synthetic data was prepared by randomly placing ∼320 modifications, calculating a map from this model and substituting it for the EM map, and using qPTxM to test how well we recover these artificial modifications.

First, we compute the correlation coefficient between the *calculated* and *experimental maps* at each unmodified residue, using 8-point interpolation at each atom position. Where the correlation coefficient is below 0.6, the local map quality is considered to be insufficient for interpretation and all possible modifications at this nucleotide are rejected (3730 out of 12664 possible modifications in the ribosome model and experimental EM map considered here). Second, we compute a set of densities at key positions in the *experimental map* that we will use to test whether there is evidence for the new atoms in the proposed modified nucleotide (Figure 2). Using tunable scalar multipliers between pairs of densities, we test whether (a) the density is strong enough at the positions of the new atoms relative to one or more adjacent reference atoms, indicating high occupancy of the new atoms, (b) the density is strong enough between the new and reference atoms compared to half a bond beyond the new atoms, indicating new atoms are more likely connected to this residue than another nearby residue, and (c) the density is strong enough at the new atoms compared to half a bond beyond, indicating a covalent bond is more likely than a nearby water molecule or ion. Modifications not passing any of these tests are also rejected (8347 out of 8934 remaining modifications here, using multipliers more stringent than the default values (see Supplementary Table S4)). Third, to select for positions where the density strongly supports both the nucleotide and the proposed new atoms, we only consider the strongest 50% *experimental map* densities at the reference atoms and the strongest 10% of *difference map* densities at the new atoms. Since these selections are based on the original, complete set of proposed modifications, this step rejects 382 of the remaining 587 modifications. Finally, qPTxM computes a score for each modification, which is calculated by taking the ratio of the *experimental map* densities at the new and reference atoms and inflating it by a factor of up to 3 for positions further from the backbone where density is expected to drop off more quickly. Modifications scoring below 0.5 are not considered further, which leaves 175 proposed modifications in the final set.

Although the true positives are some of the highest scoring candidates (Supplementary Table S5), the top candidates are residues that have never been detected as modified in *E. coli* 23S rRNA by sensitive mass spectrometry methods (33). This result suggests that although the map quality is high enough to confirm known modified residues and to guide their refinement, it is not yet sufficient to unambiguously determine the modification state of bases. Thus, a major caveat in automatic map interpretation by qPTxM is a high false positive rate, an attribute we suggest would be shared by a blinded, manual interpretation of the density map.

When examining a high-resolution structure without prior knowledge about the modifications on any of the nucleotides, qPTxM can alternatively reduce the labor required to search for possible modifications by supplying the user with a limited number of candidate sites. The researcher’s independent examination of the candidate sites is still necessary to identify cases where density can be otherwise explained, and to aid in this final step, qPTxM builds the suggested modifications onto a copy of the model for visual examination in Coot. This is assisted by a custom plugin that lets the user navigate directly to each identified location (Supplementary Figure S2). The user then manually edits the human-readable file ptms.out to remove any false positives and then re-runs qPTxM with this file to produce a molecular model with only the high-confidence modifications.

### Nucleotide conformations in the ribosome

RNA nucleobases can adopt two major conformations, *syn* and *anti*, that differ in the position of the base with respect to the sugar ring, and are interchangeable through rotation around the glycosidic bond. While the *anti* conformation is energetically more favorable than the *syn* conformation, the *syn* state affords a more compact form of the nucleotide aiding in the stability of RNA (34, 35). Detailed analysis of several RNAs and RNA-containing complexes including ribosomes indicated that most *syn* nucleobases participate in tertiary stacking and tertiary base-pairing (34). Consequently, this conformation often occurs in the functionally important regions of RNA, where *syn* pyrimidines are rarer than *syn* purines, as pyrimidine derivatives have a larger energetic penalty to adopting this conformation. The importance of *syn* conformations to RNA function is still not completely understood, and high-resolution structures are instrumental in identifying conserved *syn* nucleotides and aiding in assessment of their functional role.

Glycosidic torsion angle (χ) for each nucleotide present in our structure was measured using X3DNA suite, an integrated software system for analysis of nucleic-acid containing structures (20). The *syn* conformation was defined using IUPAC designation with a χ-angle in a range of 0° ± 90° (36). In the 23S rRNA, we observed 77 *syn* purines and 11 *syn* pyridines with good densities (Figure 4, Supplementary Table S6 and Supplementary Table S7). Following the study by Sokolski and coworkers, we divided *syn* nucleotides into two subcategories: full *syn* nucleotides, which have χ-angle between −45° and +90°, and intermediate *syn*, which have χ-angle between −90° and −45° (34). For full *syn* nucleotides the base is oriented directly above the sugar, while in the intermediate *syn* conformation the base is partially over the sugar. As a result each of these subcategories has specific functional properties. For example, the majority of full *syn* nucleotides tend to participate in tertiary stacking or tertiary base-pairing instead of secondary base-pairing (34). In our structure, *syn* nucleotides are observed throughout the 23S rRNA (Figure 4), however when examining each domain, the largest number of full *syn* nucleotides are found in the domain V of 23S rRNA, whose central loop lines the PTC and NPET (Figure 4, Supplementary Table S6 and Supplementary Table S7). One example is highly conserved nucleotide G2576, which occupies full *syn* conformation extending the stacking between G2576 and G2505. This stacking supports the single stranded rRNA segment of the central loop of domain V, which lines the NPET and is a binding site for several antibiotics (Supplementary Figure S4A). Additionally, in the PTC region, nucleotide A2503 methylated at the C2 position (m^2^A) also adopts the full *syn* conformation. It has been proposed that the presence of this methylated nucleotide in the *syn* conformation extends the stacking between A2059 and A2503, stabilizing the fold of the two single-stranded rRNA sections (13). Full *syn* nucleotides also support tertiary structure through tertiary base-pairing. For example, in the helix H20 of domain I, A330 assumes the *syn* conformation to form *trans* Watson-Crick/sugar edge A-G base pair (37) with G307, a nucleotide part of the helix H19 (Supplementary Figure S4B). This base pairing stabilizes the interaction between two helices, an interaction which is additionally facilitated by the ribosomal protein L24. Protein L24 is an assembly initiator protein and organizer of rRNA folding (38).

**Figure 4.**
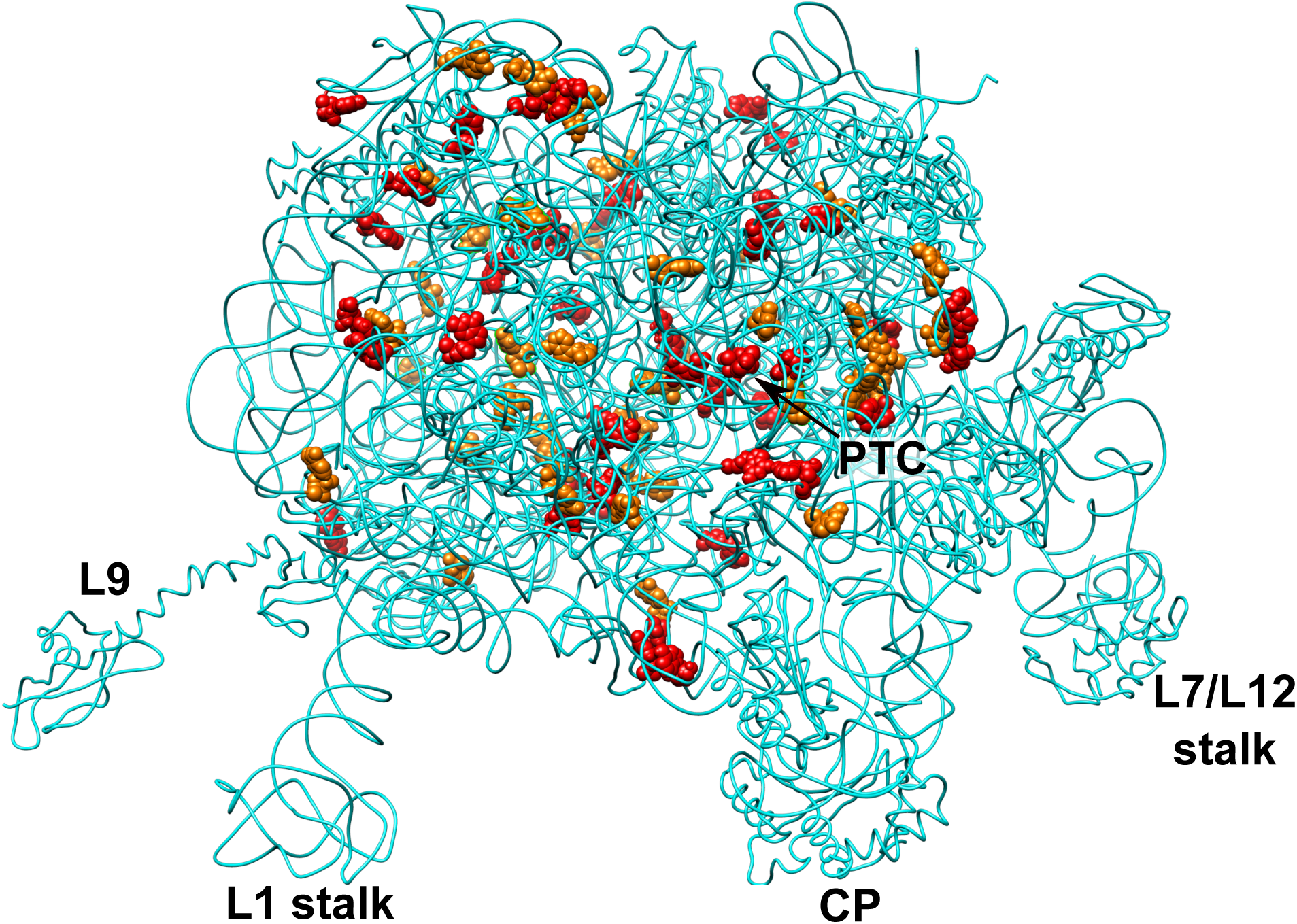
*Syn* nucleotides with good density present in cryo-EM structure are shown as spheres. Full *syn* nucleotides (90°≥ χ ≥ −45°) are shown in red, while intermediate *syn* nucleotides (−45°> χ ≥ −90°) are presented in orange. Features in the 50S subunit include the central protuberance (CP), L1 arm (L1 stalk), protein L9 (L9), L7-L12 region (L7/12 stalk), and the peptidyl transferase center (PTC).

### Solvation of the PTC and NPET

Most clinically relevant antibiotics that target the 50S subunit bind in the PTC or in the NPET region. These antibiotics inhibit protein synthesis by either perturbing the binding of tRNA at the A- or P-sites or by preventing the passage of the nascent peptide chain through the ribosomal tunnel (7). In the previous X-ray crystal structures, ordered water molecules were observed in the known antibiotic-binding sites, including PTC and NPET (14). High-resolution of the new cryo-EM structure allowed for the modelling of water molecules which were assigned using a utility supplied within Coot (19). However, while water molecules coordinating magnesium ions were evident and well-resolved throughout the structure (Supplementary Figure S5), only a few ordered water molecules were observed in the PTC and NPET. We hypothesize that water molecules and ions are diffused throughout the NPET, and could become ordered in the presence of the nascent peptide or an antibiotic.

## DISCUSSION

In the current study, we have harnessed several technological advances to obtain the highest resolution cryo-EM ribosome structure to date at an average resolution of 2.2Å. This resolution allowed for the modelling of known RNA modifications. Additionally, we present a new program for assessing the confidence with which RNA modifications can be assigned, named qPTxM. Since post-transcriptional modifications are first identified based on the properties of the map using a uniform set of criteria and subsequently examined visually by the researcher, our approach has the potential to minimize misassignment of RNA modifications in high-resolution structures, a challenge that has been encountered in the past (17, 18). For ribosomes where modifications are well known, such as the *E. coli* ribosome, the enrichment of true positives in high scoring residues can be used to assess the quality of the map.

One limitation of qPTxM is the reliance on difference maps calculated between the experimental and calculated maps, which makes the method sensitive to local resolution and B-factor refinement. A mismatch between calculated and experimental map resolutions at any given position produces errant difference density features that can hinder identification of real modifications. This is partially addressed by a filter for correlation coefficient between the experimental and calculated map at each nucleotide, and we plan to implement local resolution estimation and resolution-dependent smoothing of the calculated map in future versions of qPTxM. In the meantime, the user may generate the calculated and difference maps themselves, use another software package to compute and filter by local resolution, and supply these maps directly, overriding the steps in qPTxM that calculate these maps. Additional tests may help to reduce the false positive rate, including examining densities beyond the four locations currently tested. Another drawback of the qPTxM validation tool is that it cannot evaluate the presence of modelled pseudouridines. Thus, pseudouridines should be modelled only when there is available functional data that supports their presence or where water-mediated contact between the N1 imino group and the rRNA phosphate backbone is observed. Similarly, qPTxM cannot evaluate any modifications involving only permutations of the existing heavy atoms, such as 2-thiocytidine (s^2^C), or only hydrogenation/dehydrogenation, such as dihydrouridine (D). While we cannot search for any features that are not evident in the maps directly, we plan to expand qPTxM to handle other types of modifications, including modifications on amino acids.

The present structure can be used for comparison to other ribosome structures. While ribosomal RNA is extensively modified across all three kingdoms of life, the nature and the location of the rRNA modifications differ between kingdoms. For example, C1962 is a 5-methylcytidine (m^5^C) in *E. coli* and *Thermus thermophilus* 23S rRNA, and unmethylated in archaea and eukaryotes (39). Presence of a methyl group on C1962 extends the stacking between G1935-C1962 within the H70-H71 helical junction. As helix H71 participates in the formation of bridge B3 with 16S rRNA, it is possible that C1942 aids in strengthening of the intersubunit contact and subsequently in maintaining the 70S stability (40). Differences in the extent of modification at this nucleotide could indicate dissimilarities in the optimization of intersubunit contact and 70S stability at distinct temperatures and in different environments.

Another interesting discovery from comparison of our structure to other ribosome structures is the extent of solvation of functional regions, such as the NPET. Water molecules are known to play both functional and structural roles, especially in charged molecules such as nucleic acids. While in our structure the water molecules appear to be disordered throughout the NPET, in the high-resolution X-ray structure several well-ordered waters are identified (14). We hypothesize that these differences in the hydration pattern between the two structures are real (not a limitation of the resolution) and likely due to differences in experimental methods. One possibility is that rigid packing in the crystal lattice likely promotes ordering of the water in the X-ray structure. Alternatively, quicker vitrification during preparation of the cryo-EM sample might have aided in capturing a more biologically relevant state (41). Ultimately, understanding the hydration pattern in the ribosome structure will be important for future computational studies, calculations of the electrostatic contribution of water molecules (42) and other ions (43), and might aid in the development of new antibiotics that target the bacterial ribosome.

Lastly, our structure also allowed us to determine the conformation of each nucleotide in the structure. We have identified 88 *syn* nucleotides, predominantly in the functionally important regions of the ribosome, where they likely have a functional or structural role. Our findings are consistent with the previous analysis that revealed the overabundance of *syn* nucleotides in the antibiotic-binding sites (44). *Syn* conformation has been proposed to be associated with the higher flexibility of rRNA in these sites, resulting in pockets that are capable of the induced fit-binding mode. Thus, the identification of the *syn* nucleotides in the high-resolution structures, and especially *syn* nucleotides conserved across kingdoms could facilitate the identification of new functional sites suitable for the design of ribosome targeting drugs.

## DATA AVAILABILITY

Atomic coordinates have been deposited in the Protein Data Bank under accession number 6PJ6, the density map has been deposited in the EMDB under accession number 20353.

## SUPPLEMENTARY INFORMATION

Supplementary Data are available at NAR Online.

## ACKNOWLEDGEMENTS

We thank the Fujimori lab members for helpful discussions and comments on the manuscript and Pavel Afonine for help with cctbx map tools. We thank the UCSF Center for Advanced CryoEM, which is supported by the National Institutes of Health [S10OD020054 and 1S10OD021741] and the Howard Hughes Medical Institute (HHMI).

