## Supplementary Information for "High-resolution cryo-electron microscopy structure of the *Escherichia coli* 50S subunit and validation of nucleotide modifications"

### SUPPLEMENTARY FIGURES AND TABLES LEGENDS

Supplementary Figure S1. Fourier shell correlation (FSC) curve for the cryo-EM map. The EM map resolution was estimated to be 2.2Å using the fold-standard criterion (FSC=0.143).

Supplementary Figure S2. A custom plugin script goto_ptms.py is written out with each run of qptm.py. Supplying this script to Coot on launch along with the original and modified models allows the user to step through all suggested sites of post-transcriptional modifications.

Supplementary Figure S3. Cryo-EM density map of modified nucleotides in the 50S subunit.

Supplementary Figure S4. Example of *syn* nucleotides participating in tertiary base stacking (A) and tertiary base pairing (B). (A) Nucleotide G2576 in 23S rRNA adopts a *syn* conformation that extends the stacking between G2576 and G2505. G2505 lines the peptide exit tunnel and interacts directly with the nascent peptide. Model of VemP nascent peptide chain (PDB 5NWY) (1) is superimposed on the *E. coli* 50S structure. (B) Nucleotide A330 assumes the *syn* conformation to form trans Watson-Crick/sugar edge A-G base pair with G307. G307 and A330 are part of the two loops in helices H19 and H20, respectively. These loops are additionally stabilized through interaction with ribosomal protein L24 (cyan color). Hydrogen bonds are indicated by black dashed lines.

Supplementary Figure S5. An example of waters coordinated to magnesium ions.

Supplementary Table S1. Data collection, processing parameters, and modeling statistics.

Supplementary Table S2. Modified nucleotides in the *E. coli* 50S subunit.

Supplementary Table S3. Solvation of pseudouridines. OP refers to the oxygen, which is part of the phosphate group.

Supplementary Table S4. Accuracy of modifications identified by qPTxM in the 23S rRNA. All tests were run with an estimated resolution of 2.2Å. Final parameters differ from defaults in a selection of 60% of the reference atom densities (compared with 50%) and adjusted thresholds for ratios of densities at a nucleotide (2 more stringent, 1 more lenient). Synthetic data were generated from the same model with a random 10% of sites modified, and a map calculated from this model was used in place of the experimental map.

Supplementary Table S5. List of nucleotides known to be modified in *E. coli* 50S subunit with their qPTM scoring percentile. Scores for the assigned modifications were determined using a calculated cryo-EM map at a 2.2Å resolution. Scores were calculated on nucleotides whose correlation coefficients between experimental and calculated cryo-EM maps were at least 0.6. Modifications were limited to the 60% of nucleotides that had the strongest experimental map density at reference atom positions and the 10% of sites with the strongest difference map density at the positions of modifications. Modifications were also filtered by three thresholds on the ratios of densities in the experimental map, $d_{far} \leq 0.6•d_{new}$, $d_{far} \leq 0.4•d_{mid}$ and $d_{ref} \leq 3.5•d_{new}$ (see Methods). Scores are scaled ratios of difference and experimental map densities at the proposed and reference atom positions, respectively, and percentiles were calculated among the 176 sites passing all tests and scoring at least 0.5. NP indicates that the modification did not meet these conditions and was not assigned a score. For the nucleotide 1915, methylation was tested starting from a modelled uridine, as pseudouridine cannot be identified from the map alone.

Supplementary Table S6. *Syn* purines with good density present in cryo-EM structure. *Syn* conformation is defined by glycosidic torsion angle ($\chi$) of 0° ± 90°. Full *syn* purines (90° $\geq \chi \geq$ -45°), where the base is directly above the sugar are shown in bold, while intermediate *syn* purines (-45°> $\chi \geq$ -90°), where the base is partially over the sugar are shown in italic.

Supplementary Table S7. *Syn* pyrimidines with good density present in cryo-EM structure. *Syn* conformation is defined by glycosidic torsion angle ($\chi$) of 0° ± 90°. Full *syn* pyrimidines ( 90° $\geq \chi \geq$ -45°), where the base is directly above the sugar are shown in bold, while intermediate *syn* pyrimidines (-45°> $\chi$ $\geq$ -90°), where the base is partially over the sugar are shown in italic.

Supplementary Figure S1.

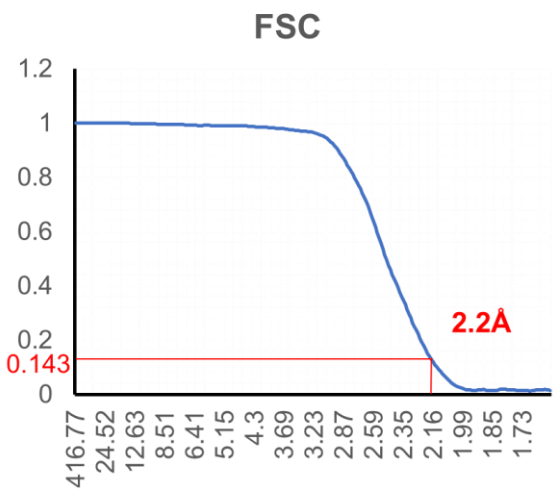

Supplementary Figure S2.

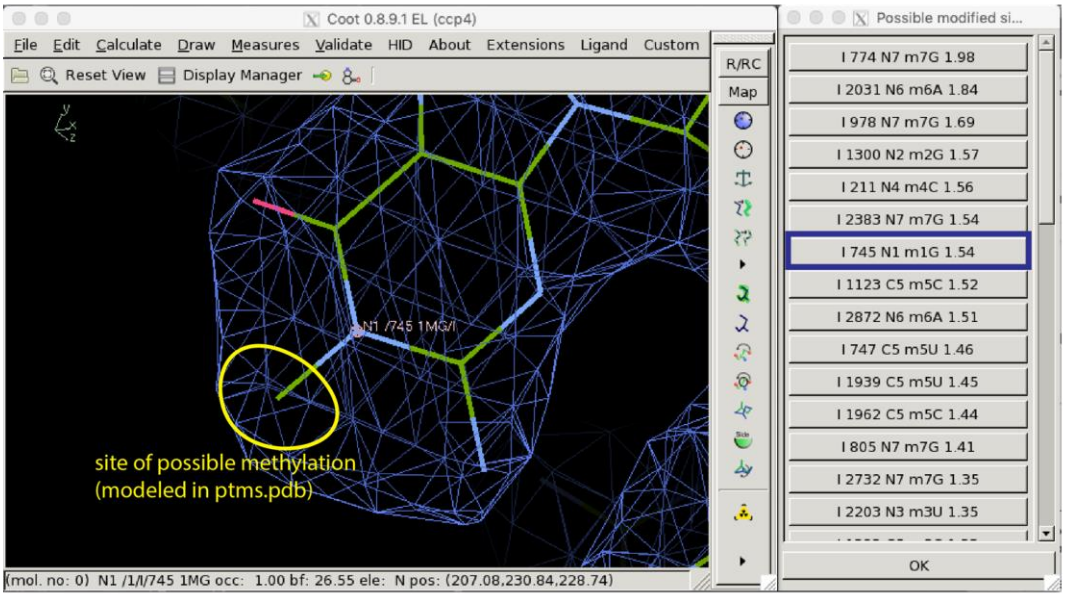

Supplementary Figure S3.

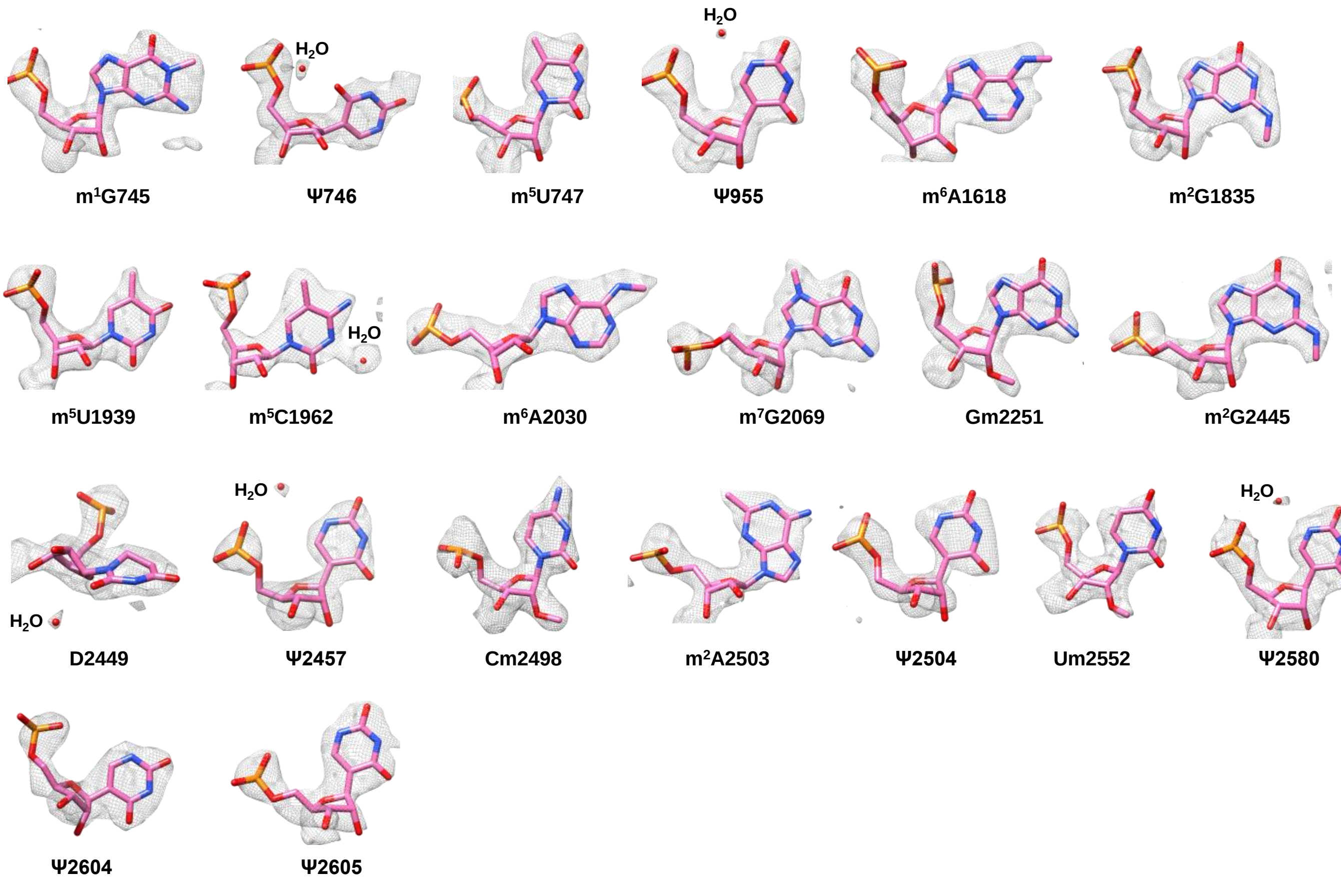

Supplementary Figure S4.

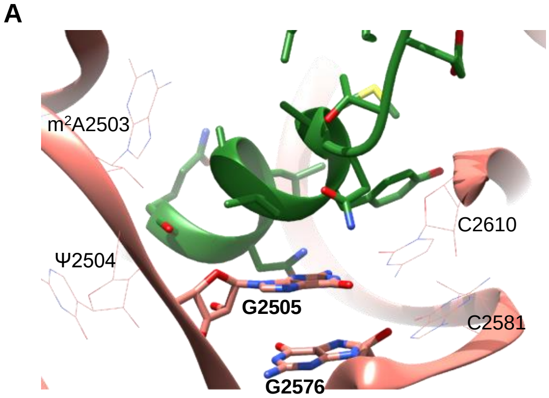


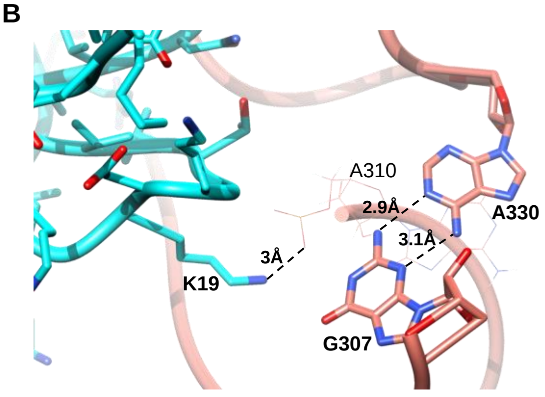

Supplementary Figure S5.

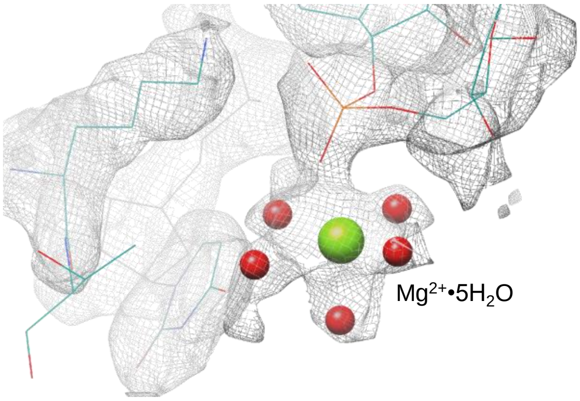

Supplementary Table S1.

| Data collection | |
|---|---|
| Electron microscope | Krios |
| Magnification | 29,000 |
| Number of micrographs | 2,688 |
| Number of particles in the map | 300,000 |
| Number of particles after classification | 144,000 |
| Pixel size (Å) | 0.822 |
| Defocus Range (µm) | -0.3 to -1.2 |
| Voltage (kV) | 300 |
| Electron dose (e-/Å$^2$) | 80 |

| Map refinement | |
|---|---|
| Resolution (Å) | 2.2 |
| Map sharpening B-factor (Å$^2$) | -70 |

| Refinement and model statistics | |
|---|---|
| Clashscore, all atoms | 4.98 |
| Protein geometry | |
| Ramachandran (%) | |
| - Favored | 95.26 |
| - Allowed | 4.68 |
| - Outliers | 0.07 |
| MolProbity score | 1.87 |
| Cβ-outliers (%) | 0.5 |
| Rotamer outliers (%) | 2.41 |
| Deviations from ideal geometry | |
| - Bonds (%) | 0.01 |
| - Angles (%) | 0.18 |
| Nucleic acid geometry | |
| Probably wrong sugar puckers (%) | 0.67 |
| Bad bonds (%) | 0.05 |
| Bad angles (%) | 0.02 |

Supplementary Table S2.

| Position | Known modification | Cryo EM density |
|---|---|---|
| 745 | $m^1G$ | + |
| 746 | Ψ | + |
| 747 | $m^5U$ | + |
| 955 | Ψ | + |
| 1618 | $m^6A$ | + |
| 1835 | $m^2G$ | + |
| 1911 | Ψ | - |
| 1915 | $m^3Ψ$ | - (a) |
| 1917 | Ψ | - |
| 1939 | $m^5U$ | + |
| 1962 | $m^5C$ | + |
| 2030 | $m^6A$ | + |
| 2069 | $m^7G$ | + |
| 2251 | Gm | + |
| 2445 | $m^2G$ | + |
| 2449 | D | + |
| 2457 | Ψ | + |
| 2498 | Cm | + |
| 2501 | $s^2C$ | - (b) |
| 2503 | $m^2A$ | + |
| 2504 | Ψ | + |
| 2552 | Um | + (c) |
| 2580 | Ψ | + |
| 2604 | Ψ | + |
| 2605 | Ψ | + |

(a) disordered region
(b) partial modification
(c) poor density for modification

Supplementary Table S3.

| **Pseudouridine** | ***Synlanti* conformation** | **$H_2O$-binding to N1** | **$H_2O$ mediated contact to** |
|---|---|---|---|
| 746 | *syn* | no | N/A |
| 955 | *anti* | yes | OP1 955<br>OP2 954 |
| 1911 | *N/A* | N/A | N/A |
| 1915 | *N/A* | N/A | N/A |
| 1917 | *N/A* | N/A | N/A |
| 2457 | *anti* | yes | OP2 2456<br>OP2 2457 |
| 2504 | *anti* | no | - |
| 2580 | *anti* | yes | OP2 2580 |
| 2604 | *anti* | no | - |
| 2605 | *anti* | no | - |

N/A = not applicable; no electron density present

Supplementary Table S4.

| Parameters | True positives | False negatives | False positives | True negatives |
|---|---|---|---|---|
| Final parameters, experimental map | 10 | 4 | 165 | 12485 |
| Default parameters, experimental map | 8 | 6 | 185 | 12465 |
| Default parameters, synthetic data | 150 | 52 | 152 | 12206 |

Supplementary Table S5.

| Nucleotide | Modification | Scoring percentile |
|---|---|---|
| 745 | $m^1G$ | 0.36 |
| 747 | $m^5U$ | NP |
| 1618 | $m^6A$ | 0.43 |
| 1835 | $m^2G$ | 0.95 |
| 1915 | $m^3\Psi$** | NP |
| 1939 | $m^5U$ | 0.17 |
| 1962 | $m^5C$ | NP |
| 2030 | $m^6A$ | 0.37 |
| 2069 | $m^7G$ | 0.01 |
| 2251 | Gm | 0.22 |
| 2445 | $m^2G$ | NP |
| 2498 | Cm | 0.24 |
| 2503 | $m^2A$ | 0.14 |
| 2552 | Um | 0.28 |

Supplementary Table S6.

| 23S rRNA | | | | | |
|---|---|---|---|---|---|
| **Domain I** | **Domain II** | **Domain III** | **Domain IV** | **Domain V** | **Domain VI** |
| **A 49** | **G 620** | *A 1286* | **G 1669** | **G 2238** | **G 2645** |
| *G 60* | *A 670* | **G 1288** | **G 1695** | **G 2250** | *G 2777* |
| **A 71** | **G 729** | **A 1301** | *G 1756* | **A 2267** | **G 2825** |
| **A 74** | *G 763* | **G 1311** | *A 1786* | **G 2286** | *A2873* |
| **A 101** | *A 764* | **G 1332** | *G 1799* | **A 2287** | **A 2879** |
| *A 118* | *G 774* | *A 1385* | **G 1929** | *G 2429* | |
| **G 177** | *G 859* | **G 1452** | *G 1930* | **A 2430** | |
| **A 196** | **A 933** | *A 1453* | **A 1936** | **A 2503** | |
| *A 199* | *G 974* | *G 1459* | *A 1981* | **A 2518** | |
| *G 301* | **A 984** | **G 1490** | | *A 2542* | |
| *A 310* | *G 1011* | *G 1568* | | **G 2576** | |
| **A 330** | **A 1021** | | | **G 2581** | |
| *G 370* | *G 1022* | | | | |
| *A 503* | *A 1128* | | | | |
| *G 506* | *A 1133* | | | | |
| **A 528** | *A 1134* | | | | |
| **A 532** | **A 1142** | | | | |
| | **G 1210** | | | | |
| | *G 1252* | | | | |

Supplementary Table S7.

| **23S rRNA** | |
|---|---|
| Nucleotide | Sugar pucker |
| **C 323** | **C2'-endo** |
| C 455 | C2'-endo |
| U 686 | C2'-endo |
| **Ψ 746** | **C1'-exo** |
| C 961 | C2'-endo |
| **U 1313** | **C4'-exo** |
| U 1394 | C3'-endo |
| C 1730 | C3'-endo |
| **U 1779** | **C4'-exo** |
| U 2609 | C2'-endo |
| **U 2884** | **C2'-endo** |
